# SYMBIO-SCAN, a full-length 16S rRNA gene-based workflow optimized for bacterial symbiont detection and discovery from long read sequencing

**DOI:** 10.64898/2026.10.26.761872

**Authors:** Carlo Croci, Giulia Iamundo, Azita Baharibabadi, Simone Sabatelli, Davide Sassera, Michele Castelli

**Affiliations:** Department of Biology and Biotechnology, University of Pavia, Pavia, Italy; Department of Entomology, Tarbiat Modares University, Tehran, Iran; Department of Biology and Biotechnologies “Charles Darwin”, Sapienza University of Rome, Roma, Italy; Fondazione IRCCS Policlinico San Matteo, Pavia, Italy; Department of Biology, University of Pisa, Pisa, Italy

**Keywords:** Coleoptera, reproductive manipulation, *Rickettsiales*, *Enterobacterales*, *Wolbachia*, Oxford Nanopore Sequencing (ONT), nutritional mutualism

## Abstract

Bacterial endosymbionts are often unculturable, making sequencing-based approaches essential for their discovery and characterization. Here we developed SYMBIO-SCAN, a dedicated workflow for recovering high-quality nearly full-length 16S rRNA gene sequences of symbionts from long reads, which provided an adequate discriminatory power for identification and for accurate phylogenetic analyses. After validation on a mock community, we applied this workflow to 30 beetle (Coleoptera) specimens from 10 species. Beetles were selected since they host a wide variety of bacterial symbionts, able to influence host physiology, ecology, population dynamics, and evolution. Despite their importance, most of these symbionts remain completely unknown or poorly characterized. By the novel approach, we obtained 80 high-quality sequences belonging to diverse bacterial lineages, mostly *Rickettsiales* and *Enterobacterales*. Putative reproductive manipulators were among the most frequently detected symbionts, including *Wolbachia*, *Rickettsia*, “*Candidatus* Lariskella”, *Arsenophonus*, and *Spiroplasma*. Moreover, three putative novel genera and one additional putative novel species were identified, including a possible *Enterobacterales* nutritional symbiont of the weevil *Holotrichapion pisi*. Overall, our results support that full-length 16S rRNA gene amplicon sequencing combined with tailored bioinformatic analyses provide a quick, cost-effective, and robust approach for investigating the diversity of host-associated bacterial symbionts, also allowing the discovery of entirely novel lineages.

## Introduction

Associations between phylogenetically diverse bacteria and eukaryotes are widespread (McFall-Ngai *et al*., 2013; Husnik *et al*., 2021), and are collectively referred to as symbioses, broadly ranging from mutualistic to parasitic (McFall-Ngai *et al*., 2013; Drew *et al*., 2021). Through long-term partnerships, bacterial symbionts can profoundly influence host physiology, ecology and evolution (Brucker and Bordenstein, 2012; Cornwallis *et al*., 2023).

In insects, the most speciose animal group (Stork, 2018), associations with bacteria are remarkably common and significantly contribute to the ecological success of their hosts (Cornwallis *et al*., 2023). Insect-associated bacterial communities are often characterized by a relatively low taxonomic complexity, with a few dominant taxa (Alberdi *et al*., 2022; Łukasik *et al*., 2026). Among these, primary symbionts have high prevalence and abundance, typically exerting relevant beneficial roles and being vertically transmitted along host generations (Douglas, 2015, 2016; Morrow *et al*., 2017; Kaltenpoth *et al*., 2025). Alongside, secondary endosymbionts are mostly facultative and characterized by more flexible transmission modes, which may facilitate host switch, often resulting in broader host ranges (Russell *et al*., 2017). Contributions of mutualistic symbionts range from nutritional complementation of specialized diets (e.g., phloem, blood or wood), provision of digestive or detoxifying enzymes, or even protection against parasites and pathogens via bioactive molecules (Cornwallis *et al*., 2023; Mondal *et al*., 2023; Kaltenpoth *et al*., 2025). On the other hand, some symbionts manipulate host reproduction, through mechanisms such as cytoplasmic incompatibility, feminization, male killing, and parthenogenesis. These strategies enhance the capability of spreading of the symbionts, and can have profound influence on their ecology and evolution, as well as on those of their hosts (Werren *et al*., 2008; Engelstädter and Hurst, 2009). Be them primary, secondary, or manipulators, symbionts are mainly intracellular, and often hosted in specialized cells, named bacteriocytes, and/or in specific body compartments (Engel and Moran, 2013; Douglas, 2015; Perlmutter and Bordenstein, 2020).

Despite their biological importance, many insect-associated bacterial symbionts are still undescribed or poorly characterized (Morrow *et al*., 2017; Wierz *et al*., 2024; Mies *et al*., 2025), particularly in the vast and neglected majority of non-model insects. Considering that symbionts are often unculturable, sequencing-based approaches are essential tools for their discovery and in-depth characterization (Handelsman, 2004). To this purpose, the 16S rRNA gene is the gold-standard marker for bacterial taxonomy and phylogeny at multiple divergence levels (Klindworth *et al*., 2013), thanks to the presence of both conserved and hypervariable regions (V1-V9). Short-read amplicon sequencing (particularly of V3-V4) is widely used for bacterial community profiling, including on insect microbiomes (Wong *et al*., 2013; Arora *et al*., 2022; Nakabachi *et al*., 2022; Moussa *et al*., 2024). Short amplicons, however, while suited for genus-level profiling based on reference databases, are very often not enough informative for robust phylogenetic analyses, which are fundamental for reconstructing evolutionary relationships among symbionts and with free-living bacteria, as well as to identify novel symbiont taxa. Long-read sequencing by PacBio or Oxford Nanopore Technology (ONT) enables the recovery of full-length 16S rRNA gene sequences, improving the resolution and accuracy of taxonomic assignments (Johnson *et al*., 2019; Matsuo *et al*., 2021). Several *in silico* tools were designed to effectively profile microbial communities from long reads (Shin *et al*., 2016; Carradec *et al*., 2020; White *et al*., 2025). On the other hand, recent improvements in long-read sequencing accuracy and costs, particularly for ONT (Agustinho *et al*., 2024), offer novel opportunities (Cuber *et al*., 2023). Accordingly, it is now possible to use ONT to obtain nearly-full length and high-quality 16S rRNA gene sequences of bacterial symbionts of insects (or other hosts) for in-depth taxonomic and phylogenetic analyses.

Here we present SYMBIO-SCAN, a workflow for obtaining high-quality sequences of bacterial symbionts from 16S amplicon ONT data through a dedicated bioinformatic pipeline, allowing their taxonomic and phylogenetic characterization. We applied this approach to a dataset of 30 Coleoptera, an ideal model for evaluating its performance and advantages, as they are the most diversified insect order, encompassing a wide range of ecological and nutritional strategies (Farrell, 1998; McKenna *et al*., 2019), and often hosting specialized bacteria (Schebeck *et al*., 2018; Salem and Kaltenpoth, 2022). Thereby, this approach allowed us to detect representatives of 36 known bacterial genera, as well as to identify three putative novel genera and one additional putative novel species.

## Material and methods

### Sample collection, DNA extraction, and host molecular characterization

A total of 30 adult Coleoptera specimens belonging to 10 species were collected from natural populations in multiple sampling locations (Supplementary Table 1). Each specimen was fixed in ethanol and stored at 4 °C until processing. Prior to extraction, to remove surface contaminants, the samples were washed in a 1% diluted bleach solution for 30 seconds, then rinsed for 30 seconds in three successive baths of deionized water (Binetruy *et al*., 2019). DNA from each individual was extracted using the NucleoSpin® Tissue kit (Macherey-Nagel, Germany), following manufacturer’s instructions. The preliminary morphological identification was verified by PCR amplification of a 710 bp fragment of the mitochondrial cytochrome c oxidase subunit I (COI) gene (Folmer *et al*., 1994), using the ExTaq polymerase (Takara Bio, Japan) and reagents, and a previously established thermal protocol (Hornok *et al*., 2016). PCR products were purified with the NucleoSpin® Gel and PCR Clean-up kit (Macherey Nagel) following manufacturer’s instructions, and Sanger sequenced in both-directions at Eurofins Genomics (Ebersberg, Germany).

### Full-length bacterial 16S rRNA gene amplification and sequencing

A PCR was performed to amplify the nearly full-length (∼1500 bp) 16S rRNA gene using the panbacterial primers 27 F (5′-AGRGTTYGATYMTGGCTCAG-3′) (Klindworth *et al*., 2013) and 1492 R (5′-CGGYTACCTTGTTACGACTT-3′) (Matsuo *et al*., 2021) each at 0.5 µM final concentration using the ExTaq polymerase (Takara Bio, Japan) and reagents. The amplification conditions were as follows: 94 °C for 3 min, 35 cycles at 94 °C for 30 sec, 55 °C for 30 sec and 72 °C for 2 min and 30 sec, plus a final elongation at 72 °C for 5 min. All PCR reactions included a negative and a positive control. PCR products were then sequenced on R10.4.1 flow cells through the ONT Lite linear/clonal amplicon sequencing service at Eurofins Genomics.

### Reads pre-processing and classification

The full bioinformatic analysis workflow was designed to process reads from each individual sample independently, and is summarized in (Supplementary Figure 1). Raw fastq reads files were subjected to quality assessment using FastQC v0.12.1 (Andrews, 2010), and NanoPlot v1.46.1 (De Coster *et al*., 2018), then quality trimmed (including adaptor removal) using fastplong v0.4.1 (Chen, 2023) with the following parameters: --length_required 500 --length_limit 2000 --trim_front 30 --cut_front --cut_front_window_size 3 --cut_front_mean_quality 25 --cut_tail --cut_tail_window_size 3 --cut_tail_mean_quality 25 --qualified_quality_phred 15 --mean_qual 20. Trimmed reads were taxonomically classified using Kraken2 v2.1.6 (Wood *et al*., 2019) with the SILVA 138.1 reference database (Quast *et al*., 2013), visualizing the results with KronaTools v2.8.1 (Ondov *et al*., 2011). Unclassified reads were discarded for downstream analyses.

### Clustering and generation of high-quality consensus sequences

The selected reads were all oriented in the 5’-3’ direction of the 16S rRNA gene with vsearch v2.30.0 (Rognes *et al*., 2016) using the SILVA reference database and converted to fasta with seqkit v2.3.0 (Shen *et al*., 2016). A first chimera filtering was performed with vsearch using the UCHIME_REF algorithm and the GOLD database as reference (Edgar *et al*., 2011). Using only >1000 bp reads (filtered with seqkit), a clustering at 97% identity was performed using vsearch with the CLUSTER_FAST algorithm, discarding clusters with <10 read size. The chimera-filtered reads were mapped to the cluster centroids using minimap2 v2.30-r1287 (Li, 2018), followed by polishing and consensus generation with racon v1.5.0 (Vaser *et al*., 2017). Consensus sequences were screened again for chimeras as described above, and, as detailed below, subjected to further steps to improve their quality by removing residual inaccuracies (e.g., small duplications, inversions, or previously unidentified chimeras). For each consensus, the closest reference sequence in SILVA was selected using BLASTn (Camacho *et al*., 2009). The respective best high-scoring segment pair (HSP) in the query-to-reference alignment was selected, assumed to represent the highest confidence core gene sequence. Then, using a custom Python script, stretches of consecutive gaps (>5 bp) with respect to the SILVA reference were identified within the terminal 200 bp at both ends of each consensus sequence, and removed together with their flanking ends. Subsequently, mismatches in the residual terminal 50 bp were identified and removed with their flanking ends. These polishing procedures allowed to balance between sequence length and accuracy.

Polished consensus sequences were subjected to a second clustering step with the CLUSTER_FAST algorithm at 97% identity, discarding clusters with a collective read size <15. For each of the remaining second-stage clusters, the respective input sequences were aligned using MAFFT L-INS-i v7.505 (Katoh and Standley, 2013), then a consensus was obtained using EMBOSS v6.6.0.0 (Rice *et al*., 2000). Each of these second-stage consensus sequences was aligned using MAFFT with its respective polished input sequences and the best BLASTn hit on SILVA, to be further refined by a custom Python script. Namely, the consensus positions differing from the SILVA reference were re-computed, using the following hierarchical criteria to distinguish genuine biological variations from putative errors. A majority rule among input sequences was always prioritized. In case of “ties”, if the same base of the SILVA reference was among the equally most abundant ones, this was preferred, except for indels. In all other cases, the first-stage read size of the input sequences was considered, and the base supported by the collectively largest read size was selected. In case of residual ties, standard IUPAC codes were used, always favoring nucleotides over gaps. Moreover, the consensus length was extended when possible. Namely, if at least one input sequence had more positions than the consensus on either end, these were added to the refined consensus, applying the criteria described above to identify the most appropriate base. The resulting refined consensus sequences were subject to a last chimera filtering, to obtain the final high-quality output sequences of the SYMBIO-SCAN pipeline, together with their classification according to the taxonomy of the best BLASTn hit on SILVA.

Here, a further a manual refinement of the results was applied based on the inspection of BLASTn results on SILVA and NCBI core_nt (accessed on August 5^th^ 2026), namely discarding potential residual chimeras, particularly those resulting from multiple chimeric junctions (less likely to be detected by UCHIME_REF), putative residual “duplicates” (defined as groups of sequences sharing ≥99.7% identity, among which the sequence with the collectively largest read size was retained), as well as non-specific eukaryotic SSU rRNA gene products (host, undigested food).

### Validation of the bioinformatic workflow on a mock community

The bioinformatic workflow described above was preliminary tested on a publicly available set of full-length 16S rRNA gene ONT reads of the ZymoBIOMICS® Microbial Community Standard sequenced on R10.4.1 flow cells (SRR23176514). The resulting consensus sequences were compared with the SILVA database and the corresponding reference 16S rRNA gene sequences available on the Zymo Research website (https://s3.amazonaws.com/zymo-files/BioPool/ZymoBIOMICS.STD.refseq.v2.zip).

### Phylogenetic analysis of nearly full-length 16S rRNA gene sequences

Phylogenetic analyses were performed on selected bacterial symbionts of interest (see Results). When multiple individuals from the same host population hosted the same bacterium, the longest sequence was taken as representative. All sequences and selected relatives downloaded from NCBI GenBank were aligned with the automatic aligner of the ARB software package v6.0.6 (Ludwig *et al*., 2004) on the 138.2 SSU Ref NR 99 ARB-SILVA database (Quast *et al*., 2013), followed by manual curation to optimize base-pairing in the predicted rRNA stems. The selected symbiont sequences were divided into three datasets according to their taxonomy, together with selected relatives and outgroup sequences, namely *Rickettsiales* (49 total sequences), *Gammaproteobacteria* (47 total sequences), and *Mollicutes* (27 total sequences). The alignment of each dataset was trimmed at both ends to the length of the shortest sequence, and a maximum-likelihood tree was inferred with phyML v3:3.3.20250515-2 (Guindon *et al*., 2010) with 100 bootstrap replicates, using the best model as indicated by jModelTest v2.1.7 (Posada, 2008) using the Akaike Information Criterion.

### Data and software availability

Sequences obtained in this work are accessible under the NCBI BioProject PRJNA1530009. Raw ONT sequencing reads were deposited to NCBI SRA under the accessions SRR40707207, SRR40994381-SRR40994409. COI host sequences (one per population) and high-quality bacterial 16S rRNA gene sequences were deposited to NCBI (COI: QB010247-QB010256, QB110673; 16S rRNA: QB021159-QB021238). The pipeline used for obtaining the high-quality 16S rRNA gene sequences is publicly available on GitHub (https://github.com/MIDIfactory/SYMBIO-SCAN).

## Results

### Molecular identification of beetles

Thirty beetle specimens from 11 populations were molecularly characterized by their COI gene sequences. This confirmed their preliminary morphological assignment to 10 different species from as many genera (including two populations of *Holotrichapion pisi*) and six families, with all specimens from each population (when available) showing 100% sequence identity (Supplementary Table 1). Specifically, most assignments were at the species-level (>99.5% best BLASTn identity on NCBI core_nt), while a single population, with one specimen available, could be identified up to the genus level, as *Onthophagus* sp. (95.5% identity to *Onthophagus pugionatus*; PQ503617).

### Validation of the SYMBIO-SCAN sequencing data analysis workflow on a mock community

Using SYMBIO-SCAN, we analysed an available package of 57,770 16S rRNA gene amplicon ONT reads from the ZymoBIOMICS® Microbial Community Standard (Supplementary Table 2). Most reads (57,511) passed quality controls, and were preliminary classified by Kraken2 in 19 genera. Roughly half of those reads (28,451) passed the initial chimera screening. After clustering at 97% identity, 83 first-stage consensus sequences were obtained, 36 of which were identified as chimeras and discarded. After polishing, the second clustering step led to obtaining 14 second-stage consensus sequences, one of which was labeled as chimeric and discarded. Finally, manual inspection of the remaining sequences and their alignments with the respective best hits on SILVA and NCBI core_nt led to discard five further chimeric sequences. Eventually, eight final high-quality consensus sequences were obtained, perfectly matching the number of bacteria present in the mock community (Supplementary Table 3). Sequences ranged from 1,290 to 1,466 bp, all showing high identity with their respective best hits on SILVA and NCBI core_nt databases (always ≥99.4%). Remarkably, they also corresponded perfectly (five sequences) or almost perfectly (three sequences with ≥99.4% identity) with reference sequences of the mock community. Therefore, we concluded that the workflow allowed a reliable discrimination between closely related bacteria (e.g., *Escherichia coli* and *Salmonella enterica* in the mock community, sharing 97.2-97.5% sequence identity depending on the 16S rRNA gene copy), as well as an accurate gene sequence reconstruction for each bacterium.

### Detection of bacterial symbionts in beetles by SYMBIO-SCAN

The full-length bacterial 16S rRNA gene was amplified from all 30 beetle specimens and sequenced using ONT, generating a total of 225,664 reads (average per sample 7,522; range 1,068-16,676) (Supplementary Table 2). For each main step of SYMBIO-SCAN, average numbers per sample and ranges obtained are reported below. After quality control (2,956 average passed reads per sample; range 498-7,298), reads were preliminary classified by Kraken2 in 10.63 average bacterial families (range 3-28) and 6.5 average genera (range 1-19). The first chimera screening led to discard 102.77 average reads per sample (range 4-411). Then, first-stage consensus sequences were obtained after clustering at 97% identity (average 9.23 per sample, range 2-19), two of which, from distinct samples, were identified as chimeras and discarded. The second clustering step led to obtaining 3.43 average second-stage consensus sequences per sample (range 1-13), for a total of 103 across the dataset, none of which was identified as chimeric by vsearch. Upon manual inspection, 23 total sequences were discarded, specifically for being chimeric (10 sequences), within-specimen duplicates (4 sequences), or eukaryotic, namely the beetle host or its plant food (one nuclear SSU rRNA sequence, two mitochondrial ones, and six chloroplastic ones). Therefore, 2.67 average high-quality sequences per sample were eventually obtained (range 1-10), for a total of 80 sequences across the whole dataset. Collectively, sequences ranged from 1,155 to 1,466 bp (average length 1,372.2 bp).

According to the hits on SILVA/NCBI core_nt and to commonly used 16S rRNA gene taxonomic thresholds (98.65-98.7% for species, and 94.5% for genus delineation) (Stackebrandt and Ebers, 2006; Kim *et al*., 2014; Yarza *et al*., 2014), the obtained sequences could be assigned to 36 described genera, 16 orders and 4 phyla (Supplementary Table 4). The identified bacteria include several primary and secondary symbionts commonly reported in association with insects, such as those belonging to the orders *Rickettsiales* and *Enterobacterales* (Table 1). Notably, three putative novel genera could be identified, as well as one possible additional novel species (see below). Insect specimens from the same population, as well as comparing the two *H. pisi* populations, were most often consistent in terms of hosted symbionts, although with some exceptions, such as *Rickettsia* in just 1/5 *H. pisi* from Guardamonte, and the differences in *Rickettsiales* symbionts among *Brassicogethes matronalis* specimens (see below). The main findings are presented below organised by bacterial taxonomy, with phylogenetic analyses performed for selected lineages.

**Table 1.**
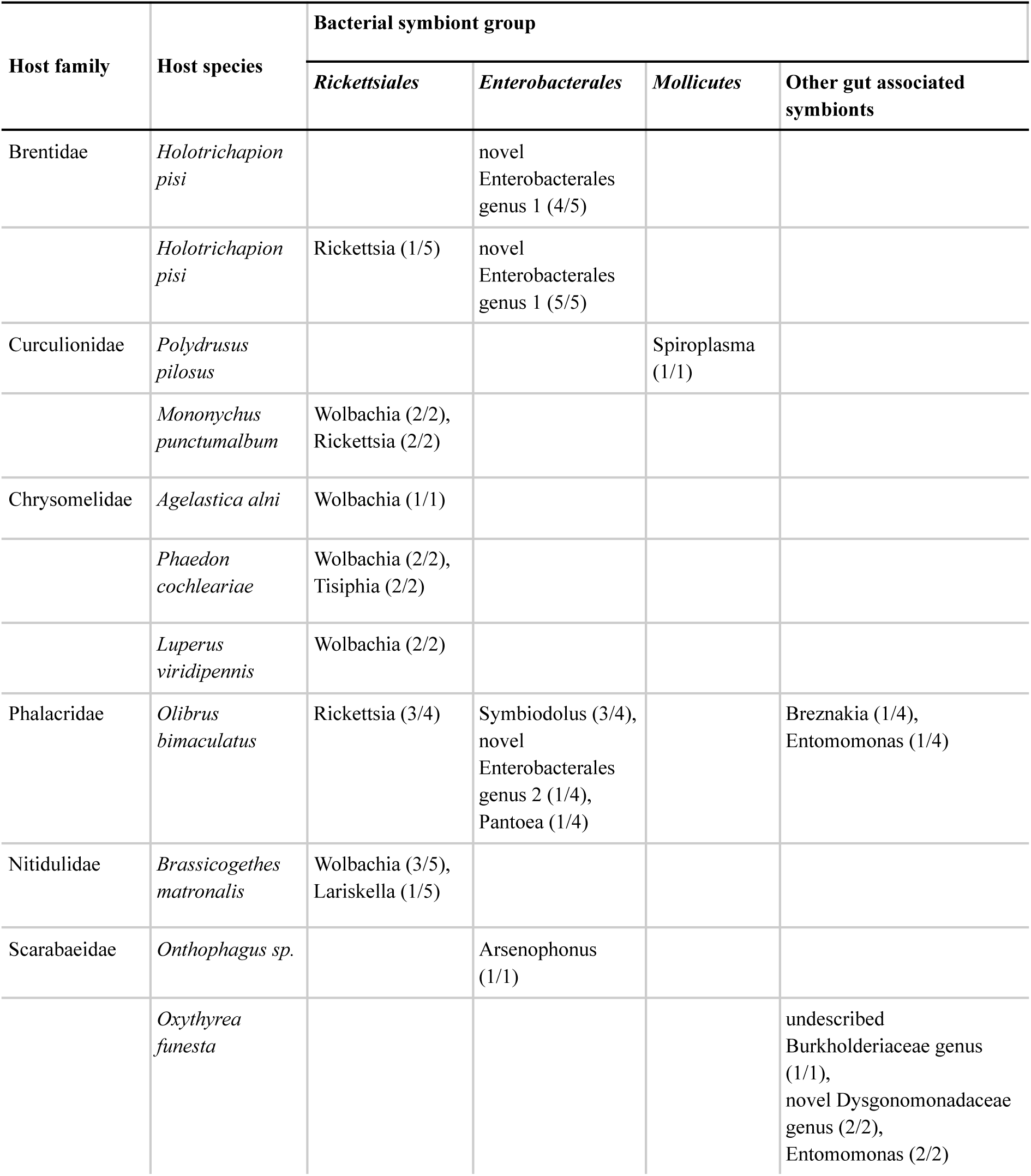
Main symbiotic bacteria identified through 16S rRNA gene amplicon sequences by SYMBIO-SCAN.

### Rickettsiales symbionts

A total of 19 high-quality sequences belonging to the order *Rickettsiales* were recovered from 15 of the 30 beetles analysed (Table 1; Figure 1), with four specimens showing coinfections involving different *Rickettsiales*, namely *Wolbachia* with either *Rickettsia* or *“Candidatus* Tisiphia” (Supplementary Table 4).

**Figure 1.**
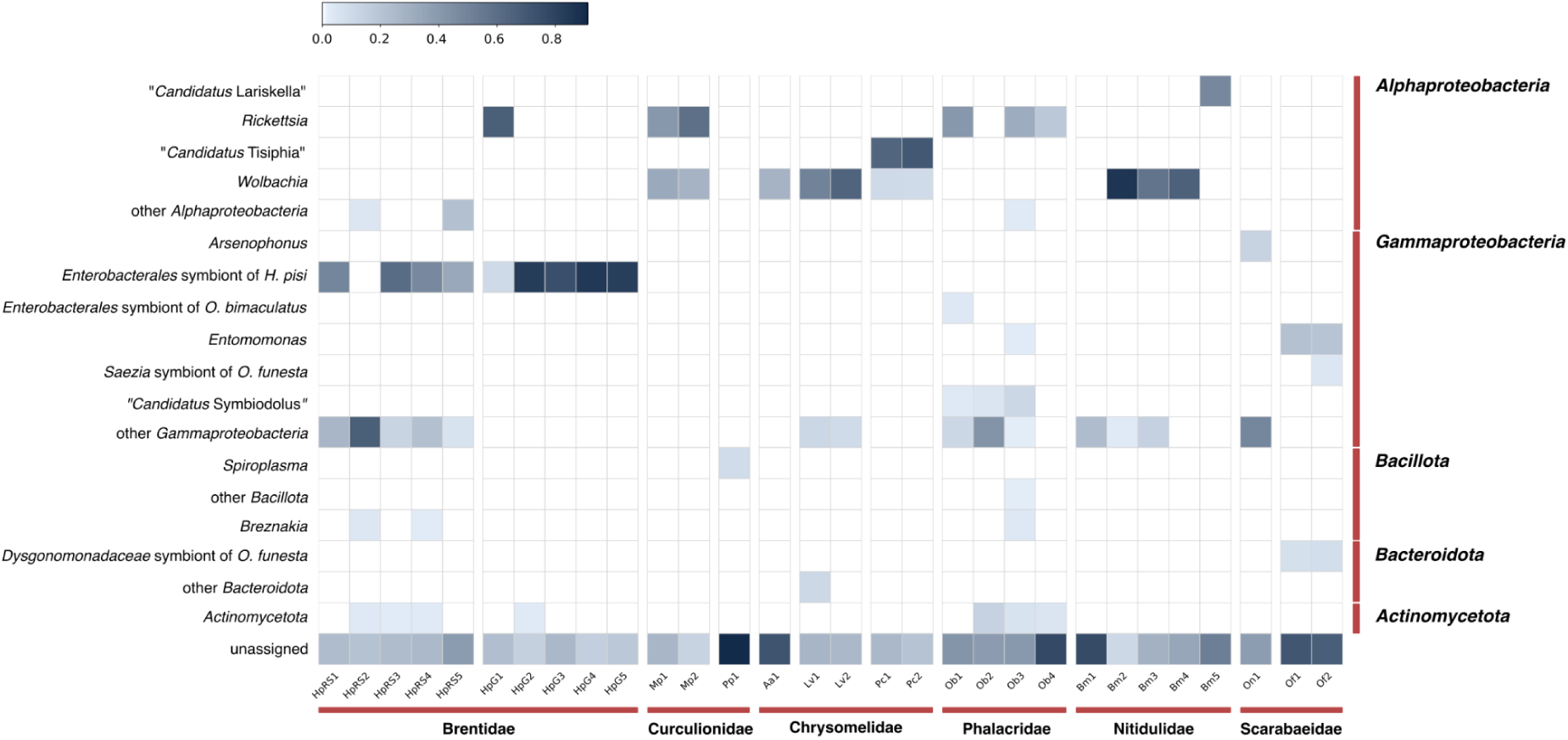
Heatmap describing the presence of the main detected symbionts and their relative abundance in the samples (as indicated by the respective read size compared to the total input reads for the clustering step for sample; see Methods).

*Wolbachia* (family *Anaplasmataceae*) was detected in multiple individuals from five different beetle species (Table 1). The newly sequenced *Wolbachia* symbionts clustered together with sequences affiliated to *Wolbachia* supergroups that are widespread in insects and other arthropods (Kajtoch and Kotásková, 2018), namely supergroup A (symbionts of the chrysomelids *Agelastica alni*, *Luperus viridipennis*, and *Phaedon cochleariae*, as well as the nitidulid *B. matronalis*) and supergroup B (symbiont of the weevil *Mononychus punctumalbum*).

“*Candidatus* Lariskella arthropodorum” (family “*Candidatus* Midichloriaceae”) was detected in a single *B. matronalis* specimen, in which, interestingly, no *Wolbachia* was detected even by the preliminary Kraken analysis. This “*Candidatus* Lariskella” symbiont clustered together with other sequences of the same genus from diverse insects (Figure 2).

**Figure 2.**
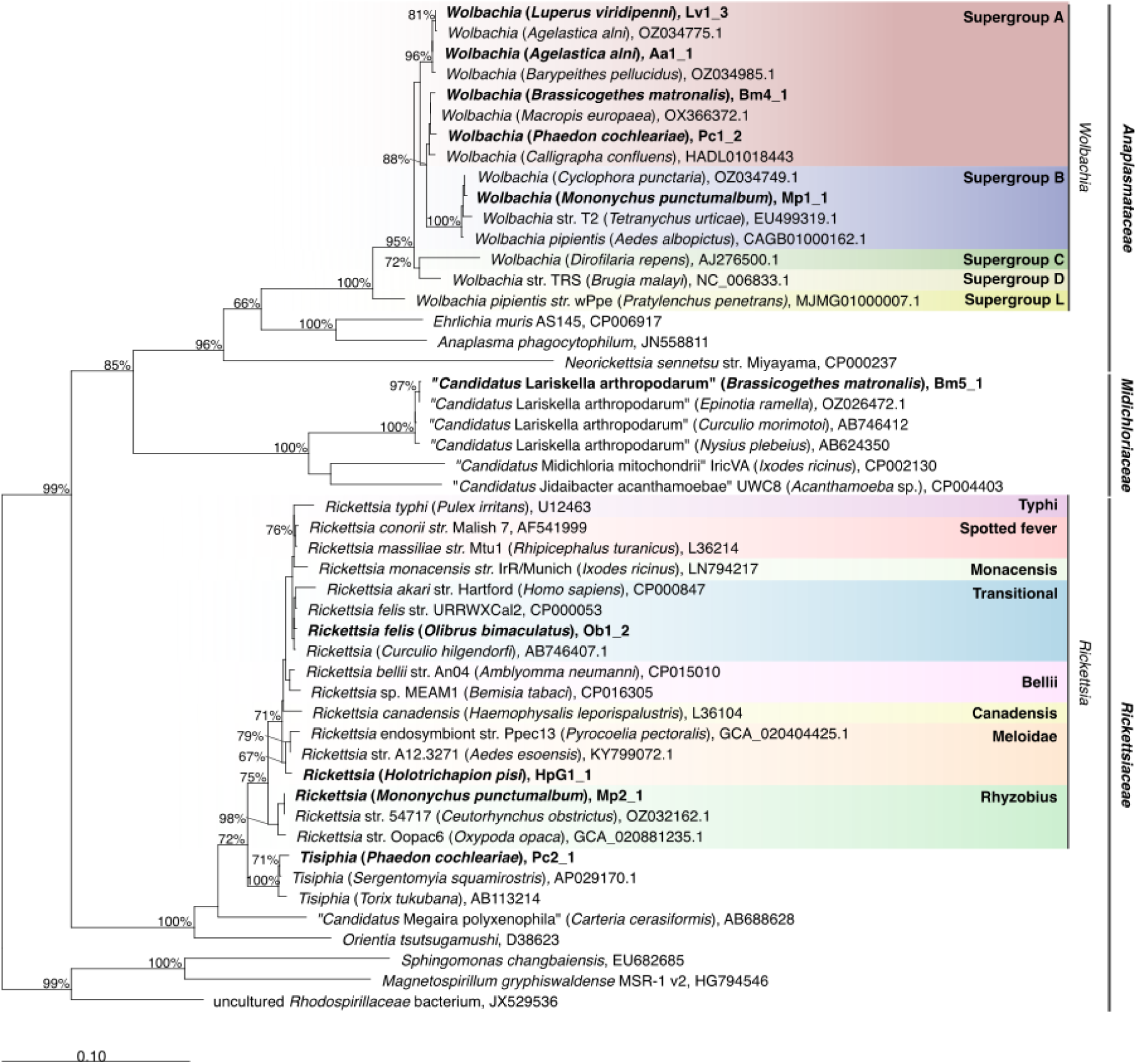
Maximum likelihood phylogenetic tree of *Rickettsiales* representatives based on 49 full-length 16S rRNA genes and 1309 sites using the model GTR+I+G. The newly sequenced symbionts of beetles in this study are evidenced in bold. *Rickettsiales* families and *Rickettsia* groups or *Wolbachia* supergroups and evidenced on the right and by colored backgrounds. Bootstrap support values are indicated at the nodes (≥65%), and the scale bar represents the estimated proportional sequence divergence.

Within the *Rickettsiaceae* family, we identified representatives of the genera *Rickettsia* and *“Candidatus* Tisiphia”. *Rickettsia* was detected in three different beetle species. The *Rickettsia* symbiont of *Olibrus bimaculatus* clustered within the *Rickettsia* Transitional group, which, alongside pathogens such as *R. felis*, also includes other symbionts of beetles, such as *Curculio hilgendorfi* (Toju *et al*., 2013). Conversely, the *Rickettsia* symbionts of *H. pisi* and of *M. punctumalbum* fell in earlier diverging positions within the genus, respectively in the Meloidae and Rhyzobius groups, alongside with symbionts of beetles and other insects. *“Candidatus* Tisiphia” was recovered only in *P. cochleariae*, and clustered with full support with other sequences of the same genus retrieved in insects and leeches (*Torix tukubana*) (Pilgrim *et al*., 2021).

### Gammaproteobacterial symbionts

A total of 18 high-quality sequences belonging to putative gammaproteobacterial symbionts were recovered from 15 of the 30 beetles analysed (Table 1; Figure 1). Many of those symbionts were affiliated to the order *Enterobacterales* (Figure 3), including representatives of two putative novel genera (16S rRNA gene sequence identity with the closest described relatives below the 94.5% genus threshold (Stackebrandt and Ebers, 2006). One novel genus was retrieved in both populations of the weevil *H. pisi*, in all but one of the ten total specimens analysed. The symbiont sequences were all identical or almost identical within and between host populations (all cross-identities ≥99.76%). In the phylogeny, sequences from the two populations form a well-supported clade, clearly separated, with a long branch, from their sister-group of the symbiont of the weevil *Sipalinus gigas*, with an identity of 86.9%. The second novel genus is represented by a sequence retrieved in one out of four specimens of phalacrid *Ol. bimaculatus*, clustering together with the primary endosymbiont of the booklouse *Liposcelis decolor* (with 86.9% sequence identity), and also showing a long branch (Figure 3). Another *Enterobacterales* symbiont detected in *Ol*. *bimaculatus* is a “*Candidatus* Symbiodolus”, clustering with full support along with sequences of “*Candidatus* Symbiodolus clandestinus” (“*Candidatus* Symbiodolus” cluster 3). The only *Onthophagus* sp. specimen hosted a bacterium clustering with high support and high sequence identity (99.9%) with the intracellular symbiont *Arsenophonus lipoptenae* (*Morganellaceae*) from the deer fly *Lipoptena*.

**Figure 3.**
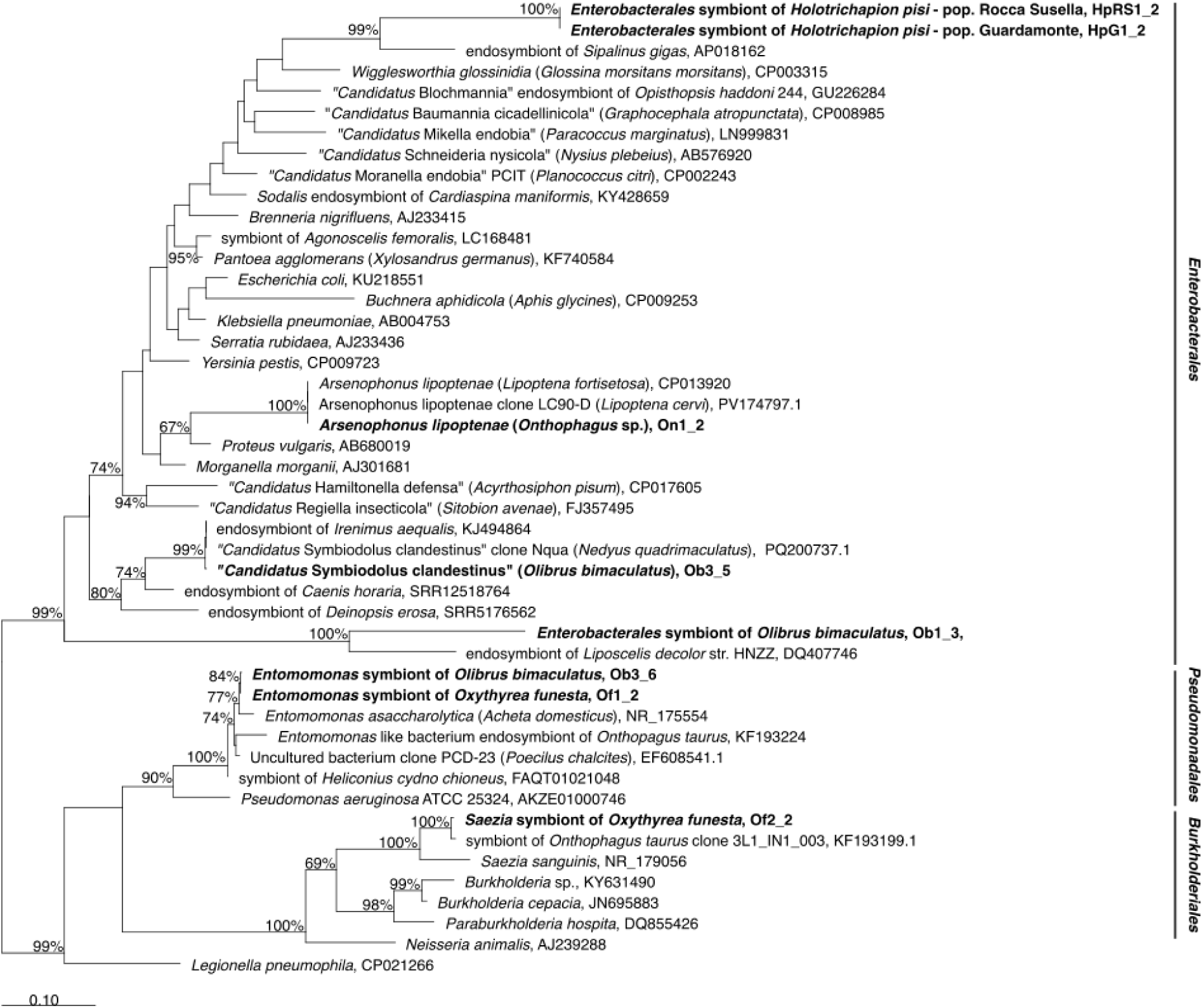
Maximum likelihood phylogenetic tree of *Gammaproteobacteria* representatives based on 47 full-length 16S rRNA gene and 1180 sites using the model TIM3+I+G. The newly sequenced symbionts of beetles in this study are evidenced in bold. *Gammaproteobacteria* orders are evidenced on the right. The tree was rooted between the family *Enterobacterales* and the other *Gammaproteobacteria* families. Bootstrap support values are indicated at the nodes (≥65%), and the scale bar represents the estimated proportional sequence divergence.

As for other gammaproteobacterial symbionts, *Entomomonas* (*Pseudomonadales*, *Pseudomonadaceae*) was found in *Ol. bimaculatus* and the scarabid *Oxythyrea funesta*. The two novel sequences cluster together within the genus, along with symbionts of the carabid *Poecilus chalcites*, the scarabid *Onthophagus taurus*, and of insects from other orders. The identity with *Entomomonas asaccharolytica* (from the house cricket *Acheta domesticus*) is ∼98.6%, which is borderline for species delimitation (threshold 98.65-98.7%) (Stackebrandt and Ebers, 2006; Kim *et al*., 2014). Moreover, a *Burkholderiales* bacterium was recovered from one of the *Ox. funesta* specimens, clustering with high support with an undescribed bacterial endosymbiont of *On. taurus* (sharing 99.4% identity), and with the closest described bacterium *Saezia sanguinis* (94.90% identity). According to species (98.65-98.7%) and genus thresholds (94.5%), this indicates that the newly sequenced bacterium and the symbiont of *On. taurus* could belong to a novel species within the *Saezia* genus (*Burkholderiales*, *Saeziaceae*).

Regarding coinfections, two *Ol*. *bimaculatus s*pecimens hosting “*Candidatus* Symbiodolus” also hosted, respectively, the novel *Enterobacterales* genus or *Entomomonas*, while an *Ox. funesta* harbored both *Entomomonas* and the undescribed *Saezia* sp.

### Other bacteria

A single high-quality *Spiroplasma* sequence (*Mycoplasmatales, Spiroplasmataceae*) was recovered from the single *Polydrusus pilosus* specimen analysed (Table 1; Figure 1). This bacterium clustered with full support with other *Spiroplasma* associated with insects, such as carabids (*Dromius quadrimaculatus* and *Clivina fossor*) or some diptera (Supplementary Figure 2).

The sequence of a *Dysgonomonadaceae* bacterium (*Bacteroidales*) was detected in two *Ox. funesta* specimens, showing very high identity (99.86-100%) with a sequence from the cockroach *Periplaneta americana* (AJ576323.1) (Egert *et al*., 2003). On the other hand, it shares a much lower identity (93.76-93.88%) with the closest hit assigned to a genus (*Dysgonomonas*) (Supplementary Table 4), indicating that the newly sequenced bacteria and the symbiont of *P. americana* could be part of the same species from an undescribed genus. Moreover, a *Breznakia pachnodae* (*Erysipelotrichales*, *Erysipelotrichaceae*) was found in *Ol*. *bimaculatus*, showing very high identity (99.64%) to a conspecific found in the scarabid *Pachnoda ephippiata* (AJ629069.1).

Most of the other high-quality sequences obtained showed high identity (above the species-threshold of 98.65-98.7%) with taxonomically and ecologically diverse bacteria, including plant-associated, environmental, and opportunistic bacteria (Supplementary Table 4). These may represent members of the insect-associated microbiota acquired through diet or the environment. Just two sequences (HpRS5_2, and HpRS2_6), while showing high identity with their respective best hits, presented lower identity to the closest taxonomically-assigned sequences, indicating that they may be part of yet undescribed species.

## Discussion

In this study, we developed SYMBIO-SCAN, a workflow for detection and accurate taxonomic classification of bacterial symbionts from host specimens, leveraging universal 16S rRNA gene PCR, ONT sequencing, and a series of optimized bioinformatic steps. Such an approach was specifically designed to obtain long, high-quality 16S rRNA gene sequences from both known and yet uncharacterized host-associated bacteria, whereas most existing ONT amplicon workflows focus on community profiling (Werren *et al*., 2008; Rodríguez-Pérez *et al*., 2021; Schacksen *et al*., 2024; Krivonos *et al*., 2025; Yergaliyev *et al*., 2025). Accordingly, particular attention was given to sequence accuracy at the individual nucleotide level, as well as to detection and removal of chimeric sequences through the integration of multiple automated and manual filtering steps. We applied SYMBIO-SCAN to a dataset of 30 specimens from ten beetle (Coleoptera) species, selected since beetles host a wide but still mostly poorly characterized diversity of bacterial symbionts (Schebeck *et al*., 2018; Salem and Kaltenpoth, 2022). This allowed us recovering 80 high-quality 16S rRNA gene sequences, including several known primary and secondary endosymbionts, as well as, strikingly, three putative novel genera and one further undescribed species, based on commonly used identity thresholds (94.5% and 98.65-98.7%, respectively) (Stackebrandt and Ebers, 2006; Yarza *et al*., 2014). Therefore, also considering the relatively small dataset size, the efficacy of the proposed workflow is established, particularly its suitability to identify previously uncharacterized host-associated bacteria.

Most beetle specimens in our dataset (20/30; Supplementary Table 4) hosted at least one known facultative symbiont, particularly including possible reproductive manipulators.

Among these manipulators, the *Rickettsiales* bacteria *Wolbachia* and *Rickettsia* were most often detected across the analysed host specimens, consistent with their known broad host ranges in Coleoptera (Kajtoch and Kotásková, 2018; Kolasa *et al*., 2018; Kajtoch *et al*., 2019). Notwithstanding known resolution limits of the 16S rRNA gene for the inner phylogeny of these two bacterial genera (Murray *et al*., 2016; X. Wang *et al*., 2020; Rodrigues *et al*., 2023), the newly sequenced symbionts clustered respectively within *Wolbachia* supergroups A and B or within *Rickettsia* Transitional, Meloidae and Rhyzobius clades, all of which are commonly associated with beetles (Kajtoch and Kotásková, 2018; Davison *et al*., 2022).

Other possible reproductive manipulators, namely *Spiroplasma*, “*Candidatus* Lariskella”, and *Arsenophonus*, were detected each in a single beetle specimen. This lower detection rate is consistent with the relatively scarce reports among Coleoptera, despite being relatively more common in other arthropods (Toju *et al*., 2013; Kolasa *et al*., 2018, 2019; Mancini *et al*., 2023; Valdivia *et al*., 2023). Similarly, the absence in our dataset of *Cardinium*, another widespread reproductive manipulator of insects, is consistent with its reportedly low distribution among Coleoptera (Kolasa *et al*., 2018). In any case, for these symbionts otherwise rare in beetles, the possible association to parasitoid wasps potentially present in the beetle specimens should be taken into account, particularly for *Arsenophonus*, often associated with parasitoids (Nováková *et al*., 2009).

It is noteworthy that “*Candidatus* Lariskella” and *Wolbachia* were found to be mutually exclusive in *B. matronalis*, consistent with another study on the same host population (Mancini *et al*., 2023) and with findings in psyllids (Morrow *et al*., 2017), while contrasting with reports of coexistence in the same host individuals of these two bacteria (Matsuura *et al*., 2012; Mulio *et al*., 2024) and of two reproductive manipulators in general (Kolasa *et al*., 2018; Xie *et al*., 2020; Guo *et al*., 2021; Wang *et al*., 2025). This could suggest a differential synergic or competitive interaction in different hosts (Rossi *et al*., 2015; Mackevicius-Dubickaja *et al*., 2025). Anyhow, the actual functional role of these potential reproductive manipulators in the analysed host specimens remains to be investigated, also considering the few available analyses from beetle hosts (Kajtoch and Kotásková, 2018; Kajtoch, 2022) compared to other arthropods, where these symbionts can act differently depending on the bacterial strain and host species, including with nutritional or defensive roles.

Other relevant facultative symbionts detected include the two recently described intracellular bacterial lineages “*Candidatus* Tisiphia” (Pilgrim *et al*., 2021; Davison *et al*., 2022) and “*Candidatus* Symbiodolus clandestinus” (Wierz *et al*., 2024), respectively in *P. cochleariae* and in *Ol. bimaculatus*. While the role in host physiology of “*Candidatus* Tisiphia” is not yet clarified, the present work further expands the known host range for this bacterium, for which currently available reports from beetles remain scarce (Küchler *et al*., 2009). Interestingly, the host specimens were collected on watercress plants (*Nasturtium officinale*) in a water canal (Supplementary Table 1), consistent with the commonly reported distribution of this bacterium in wet or aquatic environments (Davison *et al*., 2023). On the other hand, “*Candidatus* Symbiodolus”, previously reported in multiple beetles, may contribute to host nutrition and tyrosine production for cuticle sclerotization according to genome analyses (Wierz *et al*., 2024). Additionally, the host interaction of detected *Breznakia* and *Entomomonas,* while commonly reported in insect guts (Tegtmeier *et al*., 2016; J. Wang *et al*., 2020; Busse *et al*., 2021; Moser *et al*., 2023), is yet unclear and would require more detailed investigations.

The three putative novel genera and the additional putative novel species found in the analysed dataset all have close relatives that are known insect-associated endosymbionts, suggesting that the newly sequenced bacteria could have a stable or transient symbiotic relationship with their respective hosts as well. Among them, the *Enterobacterales* symbiont of the weevil *H. pisi* is noteworthy. Weevils commonly rely on bacteriome-associated primary symbiotic bacteria providing essential nutritional compounds during specific developmental stages, including tyrosine required for cuticle melanization and sclerotization (Vigneron *et al*., 2014; Mizutani *et al*., 2025). Such nutritional symbioses have evolved repeatedly in Curculionoidea, involving both ancient, co-speciating symbionts such as *Nardonella* and more recently acquired bacteria, including *Sodalis*-allied symbionts, *Curculioniphilus* (Conord *et al*., 2008; Toju *et al*., 2010, 2013)), and the unnamed symbiont of the *S. gigas* (Anbutsu *et al*., 2017), which is the closest relative of the *H. pisi* symbiont. Therefore, accounting also for its almost full prevalence in two *H. pisi* populations sampled in two different years and the absence of other known symbionts in the same specimens, it seems plausible that the newly sequenced bacterium may represent a primary endosymbiont of *H. pisi*. Contrarily, the novel *Enterobacterales* symbiont found in *Ol. bimaculatus* suggests a more transient association, being present only in one single specimen. In any case, the close phylogenetic relationship with “*Candidatus* Symbiodolus clandestinus” may suggest a facultative beneficial role in host physiology as well. The third novel genus, a *Dysgonomonadaceae* bacterium found in *Ox. funesta*, is closely related to *Dysgonomonas*, an insect-associated bacterium common in insect guts (Takahashi *et al*., 2026), being particularly abundant in the hindguts of wood-feeding termites and cockroaches, where it produces carbohydrate-active enzymes (CAZymes) to break down tough plant lignocellulose. Similarly, the genus *Saezia*, to which the putative novel species would be affiliated, is being increasingly detected in the gut and fecal communities of various insects and arthropods (Tang *et al*., 2025; Koch *et al*., 2026), including another scarbid beetle (*On. taurus*) (Estes *et al*., 2013). Interestingly, in dung beetles (Onthophagini) *Saezia* bacteria were found to possess amino acid biosynthesis pathways absent in the host genome (Jones *et al*., 2025).

In conclusion, this study illustrates the suitability of the SYMBIO-SCAN workflow for retrieving high-quality, nearly full-length 16S rRNA gene sequences using ONT for the detection and taxonomic identification of host-associated bacteria, including both known and novel ones. The workflow, herein tested to a beetle dataset, can be applied to other hosts, including but not limited to other insects and arthropods, being particularly useful for exploratory screening of poorly characterized host–microbe associations, for which reference databases are often incomplete. SYMBIO-SCAN can be used as a cheap and quick screening tool, to be followed by further genomic, imaging, and ecological investigations to reveal the actual nature of the association of the detected bacteria with their hosts. Overall, improved approaches for characterizing arthropod-associated bacteria are crucial for a better understanding of host–microbe interactions, being instrumental for the management of agricultural pests and vectors of plant, livestock, and human pathogens within a One Health approach.

## Acknowledgements

This project was funded under the National Recovery and Resilience Plan (NRRP), Mission 4 Component 2 Investment 1.4—Call for tender No. 3138 of 16 December 2021, rectified by Decree n.3175 of 18 December 2021 of the Italian Ministry of University and Research funded by the European Union—NextGenerationEU to DS. The authors wish to thank Sophie Melis and Lorenzo Losa for beetle sampling, Paolo Losi for assistance in laboratory molecular biology procedures, and Roberto Marangoni for feedback on the pipeline and manuscript.

## Supplementary figures and tables captions

**Supplementary Figure 1** - Graphical representation of the bioinformatic workflow used to obtain high-quality sequences using Oxford Nanopore Sequencing.

**Supplementary Figure 2.** Maximum likelihood phylogenetic tree of *Mycoplasmatales* representatives based on 27 full-length 16S rRNA genes and 1373 sites using the model GTR+I+G. The newly sequenced symbionts of beetles in this study are evidenced in bold. Bootstrap support values are indicated at the nodes (≥65%), and the scale bar represents the estimated proportional sequence divergence.

**Supplementary Table 1** - Sampling metadata for the beetles individuals of 10 species representing ten genera within Coleoptera order

**Supplementary Table 2** - number of reads, OTUs, consensus sequences, and chimera trimmed and generated in the workflow

**Supplementary Table 3** - Sequences length, percentage identity and taxonomical classification of the obtained high-quality consensus sequences of the ZymoBIOMICS® Microbial Community Standard reads (sample SRR23176514)

**Supplementary Table 4** - Sequence length, percentage identity of the obtained high quality sequences from beetle specimens and their inferred taxonomic classification

## Notes

https://github.com/MIDIfactory/SYMBIO-SCAN/tree/main

### Competing Interest Statement

The authors have declared no competing interest.

