## Supplementary figures and images for "SYMBIO-SCAN, a full-length 16S rRNA gene-based workflow optimized for bacterial symbiont detection and discovery from long read sequencing"

### Supplementary Figure 1.pdf

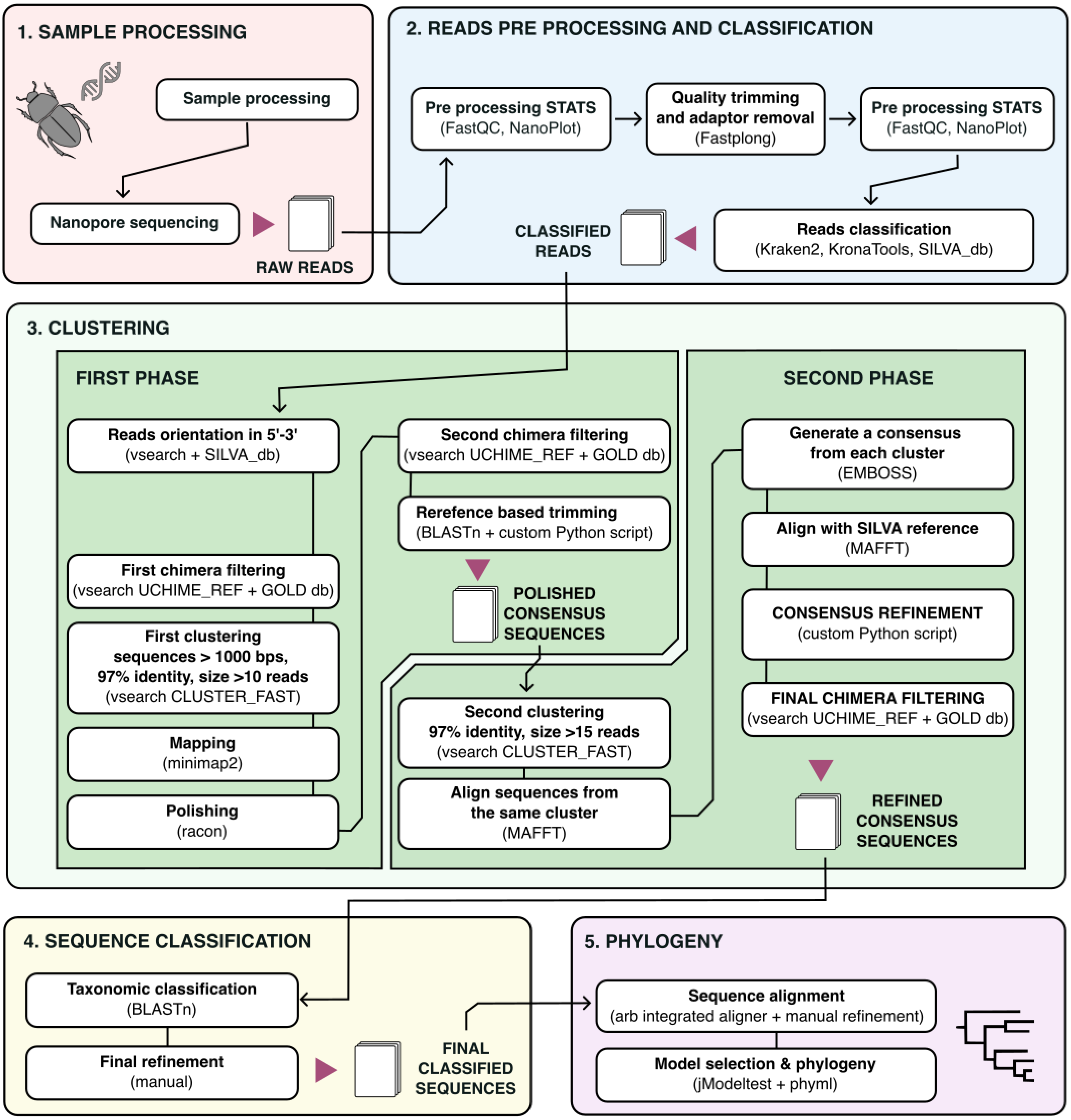

### Supplementary Figure 2.pdf

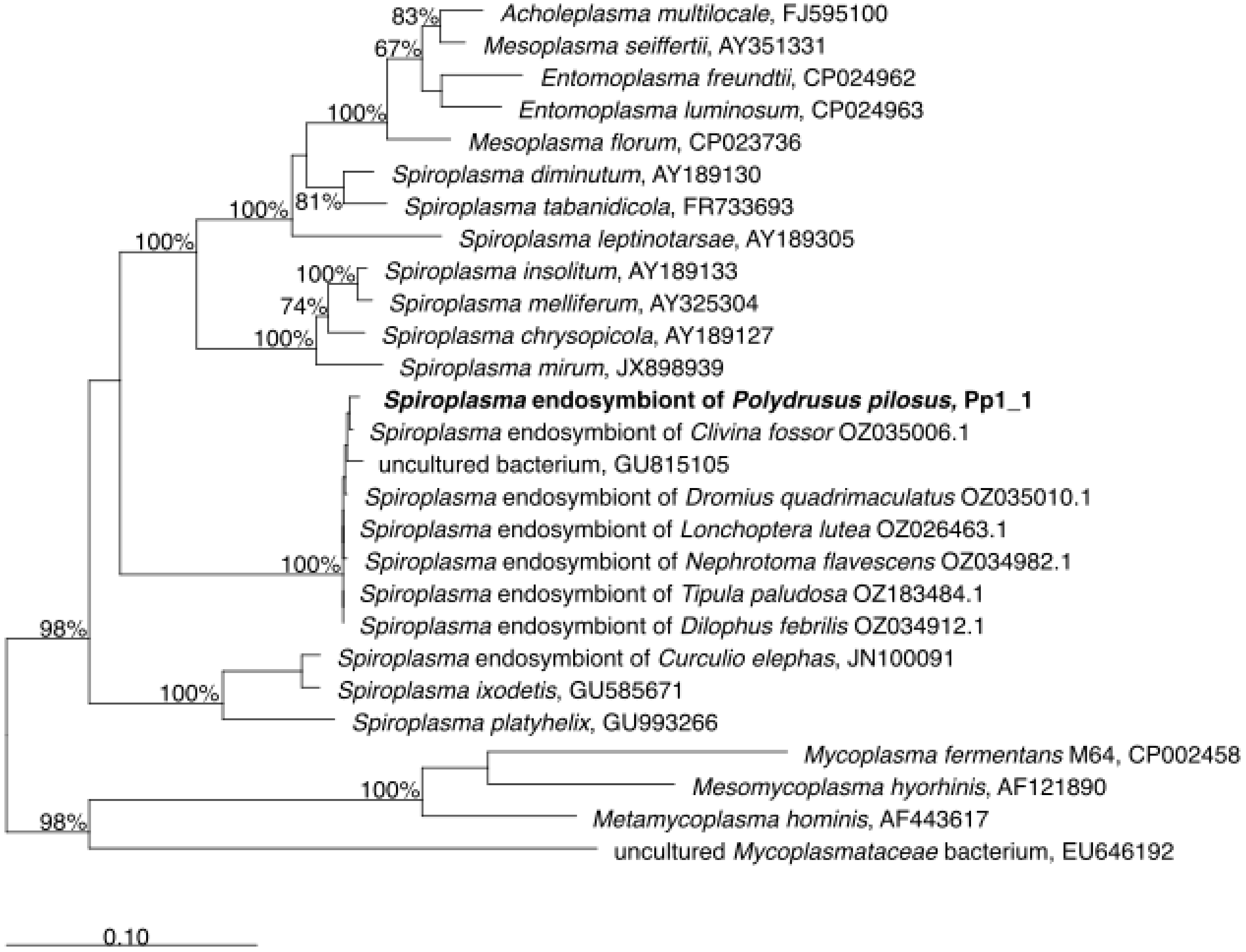
